# OAC-39 is an *O*-acyltransferase required for the synthesis of maradolipids in the dauer larva of *C. elegans*

**DOI:** 10.1101/2021.10.15.464527

**Authors:** Sider Penkov

**Affiliations:** Institute for Clinical Chemistry and Laboratory Medicine, University Clinic and Faculty of Medicine, TU Dresden, Dresden, Germany; Center for Membrane Biochemistry and Lipid Research, University Clinic and Faculty of Medicine, TU Dresden, Dresden, Germany

## Abstract

Upon overcrowding or low food availability, the nematode *C. elegans* enters a specialized diapause stage for survival, called the dauer larva. The growth-arrested, non-feeding dauer larva undergoes a profound metabolic and physiologic switch underlying its extraordinary stress resistance and longevity. One of the metabolic signatures of dauer larvae is the accumulation of the disaccharide trehalose, which lowers the sensitivity of worms to desiccation and hyperosmotic shock. Previously, we have found that trehalose is incorporated as a headgroup into dauer-specific 6,6’-di-*O*-acyltrehalose lipids, named maradolipids. Despite comprising a bulk fraction of the polar lipids in dauer larvae, little is known about the physiological function of maradolipds because the enzyme(s) involved in their synthesis has not yet been identified. Here, we report that the dauer-upregulated *O*-acyltransferase homolog OAC-39 is essential for the synthesis of maradolipids. This enzyme is enriched at the apical region of the intestinal cells of dauer larvae, where it might participate in the structuring of the gut lumen. As OAC-39 is most probably responsible for the last step of maradolipid synthesis, its identification will pave the way for the elucidation of the function of this obscure class of lipids.

## Introduction

A universal response of organisms to harsh environmental conditions is the cessation of growth and the entry of a prolonged metabolically repressed state. This phenomenon is widespread in the animal kingdom, too, as diverse metazoan species possess the ability to undergo programmed and reversible developmental arrest termed diapause (1). Studying diapause has been proven useful for understanding the signaling cascades and molecular mechanisms of controlled transition to quiescence at the organismal level, as well as the emergence of stress resistance (1).

Among the well-known examples of diapause is the dauer larva of the roundworm *C. elegans* (from German “dauer” - enduring) (2, 3). The dauer program is activated in response to increased population density, low food availability, and high ambient temperature and is regulated by various conserved signaling pathways such as the guanylate cyclase, TGFβ, insulin, and a steroid hormone pathway (2–4). In postembryonic development, there are four larval stages preceding the formation of adult *C. elegans* worms, separated by molting events (5). Dauer larvae form as an alternative third larval stage and are metabolically and physiologically distinct from all other life stages of *C. elegans*. They are sealed off from the surrounding environment by a tight cuticle limiting chemical exchange and providing resistance to oxidants, detergents, etc. (2, 3, 6, 7). This sealing extends to the buccal opening, thus preventing not only the infusion of dangerous chemicals but also food intake (2, 3, 8).

Since dauer larvae do not feed, they rely on stored energy reserves, mostly consisting of fat droplets, for energy production and prolonged survival (2, 3). To maximize the utilization of these reserves, dauers undergo a profound metabolic switch (2, 3, 9–12). This is manifested in general suppression of the anabolism and the pathways that support it such as TCA cycle and oxidative phosphorylation (2, 9–12). Instead, dauers very slowly catabolize the fat reserves to power carbohydrate production via the glyoxylate shunt and gluconeogenesis (2, 9–12). The resulting sugars then maintain the energy production through aerobic glycolysis (2, 9–12). The main product of gluconeogenesis is the disaccharide trehalose, which accumulates in large amounts in dauer larvae (2, 9, 10, 13). Interestingly, apart from providing a substrate for glycolysis and energy production, trehalose synthesis has been implicated in various other aspects of dauer physiology. First, we have shown that it regulates the redox state and the aforementioned steroid hormone signaling that governs dauer formation (14). Second, trehalose itself acts as a stress protectant molecule that supports the higher resistance of dauers to desiccation and hyperosmotic stress (15, 16).

Apart from its autonomous functions as a free disaccharide, we have discovered that trehalose is also incorporated as a headgroup into two related classes of dauer-specific lipids (17, 18). The more abundant class is comprised of diacyltrehaloses that we named “maradolipids”, formed by esterification of trehalose at 6- and 6’-position with two fatty acids (also known as *O*-acylation) (17). Less represented is the class of monoacyltrehaloses or “lyso-maradolipids” in which trehalose is singly acylated (18). As described by us and later confirmed by other groups, both classes are present only in dauer animals and are not detectable in other developmental stages (17–19). A shared feature of both classes is a specific fatty acid composition. They show a pronounced enrichment of monomethyl branched-chain fatty acids (mmBCFAs) (17–19), which reaches up to approximately 66 mol% in maradolipids (17).

The trehalose lipids constitute a significant portion of the polar lipids in dauer larvae: we have estimated that for every seventeen molecules of membrane phospholipids, there should be about one maradolipid molecule (17). Despite their abundance, however, still little is known about the functions they perform. Indirect evidence has been collected through genetic ablation targeting the production of the trehalose headgroup or the mmBCFAs, the two pathways that provide the primary building blocks for maradolipid synthesis (**Figure 1A**) (17). These experiments have shown that trehalose and mmBCFAs are necessary for the proper structuring of the intestinal brush border (17). Specifically, electron microscopy showed that blocked or diminished production of either entity resulted in the reduction or, in some cases, the complete disappearance of a coat-like electron-dense material that covers the luminal apical surface and the microvilli of dauer intestinal cells (17). This implies that maradolipids could be involved in forming this coat-like structural entity, which, in turn, might contribute to the chemical isolation of dauer larvae from the external environment. However, a direct examination of the functions of (lyso-)maradolipids requires the knowledge of what enzyme(s) catalyzes the esterification of trehalose to fatty acids. So far, such a protein has not been identified.

**Figure 1.**
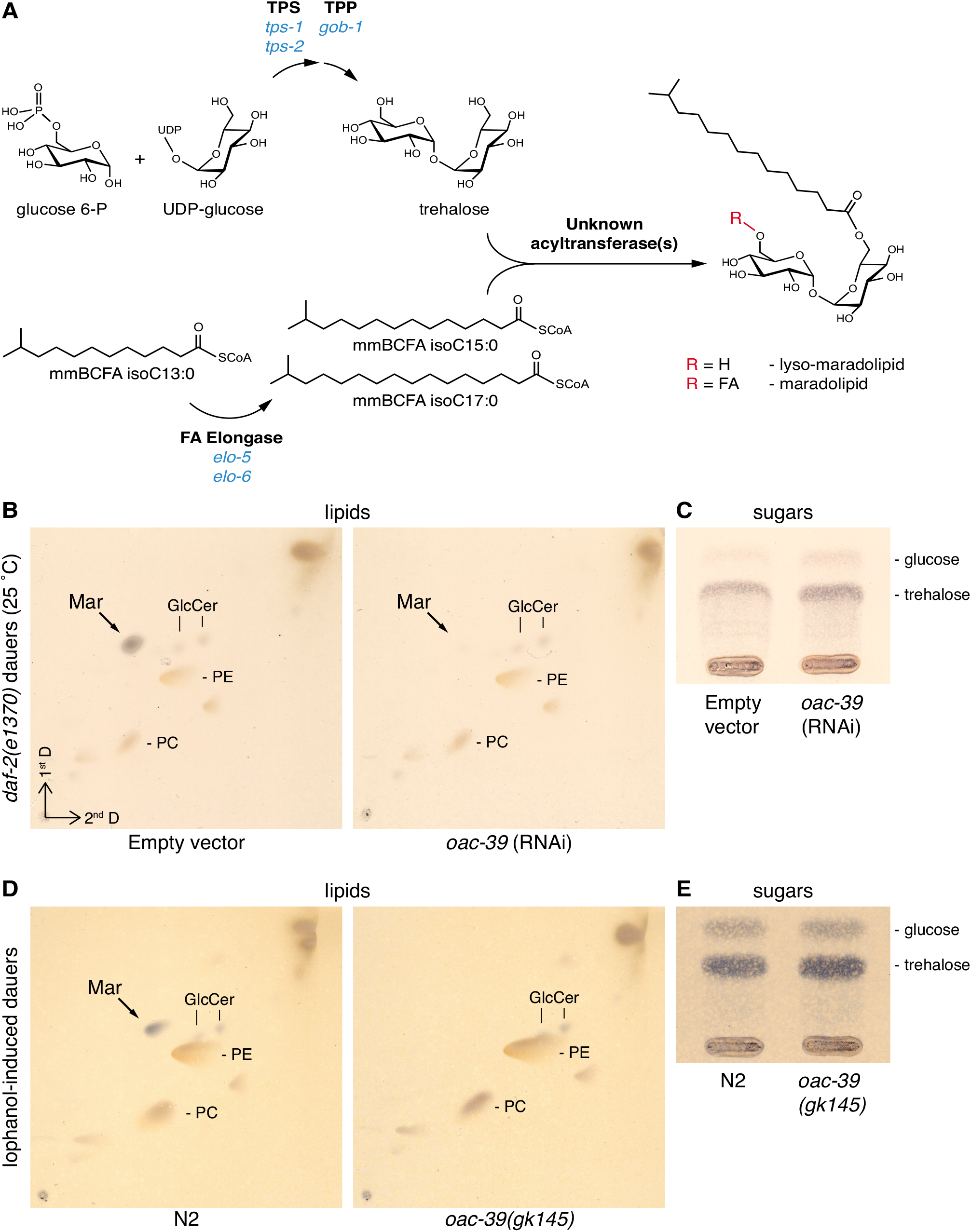
Genetic ablation of OAC-39 leads to depletion of maradolipids but not of the trehalose headgroup in dauer larvae. **A)** Pathway of (lyso-)maradolipid synthesis. The trehalose headgroup is synthesized in two steps by conjugation of glucose 6-P and UDP-glucose by trehalose 6-P synthase (TPS, encoded by two *C. elegans* genes, *tps-1* and *tps-2*) and subsequent dephosphorylation by trehalose 6-P phosphatase (TPP, encoded by *gob-1*). The major mmBCFAs in *C. elegans*, isoC15:0 and isoC17:0, are derived from a shorter precursor isoC13:0 via elongation performed by fatty acid elongase (encoded by *elo-5* and *elo-6*). The esterification of fatty acids to trehalose is performed by a yet unknown acyltransferase(s), most probably via a transfer from an acyl-CoA conjugate. Depicted is one of the most common products, a (lyso-)maradolipid containing a isoC15:0 moiety. While (lyso-)maradolipds are enriched in mmBCFAs, other fatty acids are also incorporated (not shown). **B)** 2D-TLC of lipids derived from *daf-2(e1370)* dauer larvae subjected to control RNAi (empty vector) or RNAi against *oac-39*. **C)** 1D-TLC of sugars from *daf-2(e1370)* dauer larvae on control RNAi or RNAi against *oac-39*. **D)** 2D-TLC of lipids from lophanol-induced wild-type (N2) or *oac-39(gk145)* mutant dauer larvae. **E)** 1D-TLC of sugars from lophanol-induced wild-type (N2) or *oac-39(gk145)* mutant dauer larvae. In panels **B)** and **D)**, Mar – maradolipids, GlcCer – glucosylceramides, PE – phosphatidylethanolamines, PC – phosphatidylcholines.

Here, we show evidence that the *O*-acyltransferase homolog OAC-39 is required for the synthesis of maradolipids. OAC-39 is dauer-specific and its expression is initiated at the onset of dauer arrest, coinciding with the appearance of maradolipids. A protein reporter shows localization in the intestinal cells with a particular apical enrichment. Hence, OAC-39 is most probably the enzyme that catalyzes the last step of maradolipid synthesis at the border to the intestinal lumen. Its identification will facilitate the future investigation of the roles of trehalose lipids in dauer larvae.

## Materials and Methods

### Chemicals and *C. elegans* strains

Lophanol was produced in the laboratory of Prof. H.-J. Knölker. All other chemicals were from Sigma-Aldrich (Taufkirchen, Germany) unless specified.

Wild-type (N2) and *oac-39(gk145) C. elegans* strains were provided by the Caenorhabditis Genetics Centre (CGC). A strain expressing OAC-39::eGFP protein reporter was generated by the Genome Engineering facility of MPI-CBG, Dresden, Germany. Shortly, a modified genomic clone WRM0629D_G05(pRedFlp-Hgr)(tag-40[13054]::S0001_pR6K_Amp_2xTY1ce_EGFP_FRT_rpsl_neo_FRT_3xFlag)dFRT::unc-119-Nat containing a copy of the *oac-39* gene (also known as *tag-40* or R02C2.3) tagged inframe at the C-terminus with a cassette containing an eGFP sequence was generated and used for biolistic transformation of *unc-119* mutant strain as previously described (20, 21). Worms bearing an extrachromosomal array of the construct and expressing the OAC-39::eGFP reporter were selected based on *unc-119* rescue and eGFP fluorescence.

### Growth of *C. elegans* strains, generation of dauer larvae, and RNAi by feeding

All *C. elegans* strains were routinely propagated on NGM-agar plates complemented with *E. coli* NA22 grown in LB medium (5). The temperature-sensitive dauer constitutive strain *daf-2(e1370)* was grown at 15°C, a temperature at which it undergoes reproductive growth (22).

To obtain dauer larvae of *daf-2(e1370)* and induce RNAi against *oac-39*, adults grown at 15°C were treated with hypochlorite solution. The resulting embryos were grown on NGM-agar plates at the restrictive temperature 25°C (22) for 72 hours on plates containing *E. coli* HT115(DE3) dsRNA-producing strains (23) purchased from the dsRNA feeding library of Source BioScience, UK.

For the preparation of dauer larvae of the wild-type (N2) and *oac-39(gk145)* strains, worms were grown on a lophanol-containing medium as described before (24). Shortly, to avoid cholesterol contamination, chloroform-extracted agarose plates were seeded with *E. coli* NA22 grown in a cholesterol-free DMEM medium. Lophanol was added to the bacteria such that the final concentration according to the total volume of the agarose was 13 μM. Embryos from hypochlorite-treated worms were grown on this medium for two consecutive generations. In the second generation, worms uniformly arrested in dauer state.

Dauer larvae were washed three times in ddH_2_O and snap-frozen in liquid nitrogen for biochemical analysis.

### Organic extraction and thin-layer chromatography (TLC)

Frozen dauer larvae were homogenized by three rounds of freezing and thawing in an ultrasonication bath and extracted using a standard method (25). After phase separation, lipids and hydrophilic metabolites were recovered from the organic and aqueous phases, respectively. Samples were normalized according to the number of worms and applied to 10 cm HPTLC silica gel 60 matrix plates (Merck, Darmstadt, Germany). For lipid analysis, organic phases were subjected to 2D-TLC with chloroform-methanol-water (45:18:3, v/v/v) as the first mobile phase and chloroform-methanol-32% ammonia (60:35:5, v/v/v) as the second. Aqueous phases were used for sugar detection via 1D-TLC using chloroform-methanol-water (4:4:1, v/v/v) as the mobile phase. Both lipids and sugars were visualized after staining with Molisch reagent.

### Fluorescent microscopy

Worms expressing the OAC-39::eGFP reporter were grown on NGM-agar plates complemented with *E. coli* NA22 until the increasing population density caused spontaneous dauer formation. Dauer larvae that showed eGFP signal were manually selected and transferred to glass slides (Thermo scientific, Superfrost Plus) with 2% agarose pads and anesthetized with 20 mM sodium azide in M9 buffer. After removing the excess liquid and mounting of 0.17 +/-0.005 mm coverslips (Menzel-Glaeser), fluorescence confocal microscopy was performed with a Zeiss Axiovert 200M scanning confocal microscope equipped with a Zeiss Plan-Apochromat 10x 0.45 and a Zeiss Plan-Neofluar 40x 1.3 oil immersion objective. eGFP was excited at 488 nm, and fluorescence was detected at the emission band of 490-540 nm. Images were handled using Fiji (26).

## Results

### OAC-39 is a dauer-specific enzyme required for the synthesis of maradolipids

We set out to identify candidate enzymes that could catalyze the esterification of fatty acids to trehalose. Trehalose lipids have been primarily associated with microorganisms such as *Mycobacterium*, *Rhodococcus*, and *Nocardia*, producing them as structural components of the cell envelope (27–29). Hence, *C. elegans* might express a homolog of some bacterial enzyme obtained through horizontal gene transfer. However, we could not find homologs of any of the known bacterial trehalose-modifying enzymes in the *C. elegans* genome. Although a homolog originating from bacteria can not be completely excluded, we hypothesized that worms have independently evolved own enzyme(s) capable of trehalose acylation. One unusually expanded gene family in *C. elegans* that can potentially carry out such functions is comprised of sixty genes encoding *<u>O</u>*-<u>ac</u>yltransferases (*oac*). Most of these genes remain functionally uncharacterized. One of the *oac* members, *oac-39* (also known as *tag-40* or R02C2.3), appeared to have a strong dauer-specific expression (Erkut and Kurzchalia, unpublished transcriptome analysis). Indeed, according to the *C. elegans* Dauer Metabolic Database (http://www.dauerdb.org), a resource dedicated to dauer gene expression patterns, *oac-39* was expressed at very high levels in dauer larvae but hardly detectable in other developmental stages (30). Moreover, *oac-39* expression was induced late in dauer formation (30), corresponding to the developmental stage at which we first detected maradolipids (17). Hence, *oac-39* encodes a dauer-specific *O*-acyltransferase that might acylate trehalose.

To address the involvement of OAC-39 in the synthesis of trehalose lipids, we tested if the loss of *oac-39* expression would affect their levels. As shown before, maradolipids are readily detectable via two-dimensional thin-layer chromatography (2D-TLC) of lipids derived from dauer larvae (17). First, we applied RNAi against *oac-39* in larvae undergoing dauer development. To have a synchronous entry into dauer state, we made use of the temperature-sensitive dauer-constitutive mutant *daf-2(e1372)* that undergoes 100% dauer arrest at the restrictive temperature of 25 °C due to the impaired function of the insulin receptor homolog DAF-2 (22). While 2D-TLC of control *daf-2* dauers treated with empty vector showed a very prominent maradolipid band (**Figure 1B**) (17), *oac-39* RNAi caused an almost full depletion of maradolipids (**Figure 1B**) but not of non-esterified free trehalose (**Figure 1C**).

It is known that RNAi can cause off-target effects against loci with sequence similarity to the primary target (23). As *oac-39* is homologous to other *oac* genes, the depletion of maradolipids upon *oac-39* RNAi could thus stem from the decreased expression of some other *oac* member rather than of *oac-39* itself. To exclude this possibility and complement the RNAi approach, we used a mutant strain, *oac-39(gk145)*, bearing a large 1855 base pairs deletion in the coding region of *oac-39*. This time, we induced dauer formation by substituting the cholesterol in the growth medium with another sterol, lophanol (24). This alternative method also yields 100% synchronous dauer larvae by inhibiting the synthesis of steroid hormones called dafachronic acids, ligands of the steroid hormone receptor DAF-12 (24, 31). Wild-type (N2) dauers grown on lophanol showed regular maradolipid content (**Figure 1D**) (17). In contrast, *oac-39* mutant dauers had no detectable maradolipids (**Figure 1D**). However, the levels of free trehalose in *oac-39(gk145)* did not decrease compared to the wild-type (**Figure 1E**). Together, these experiments show that OAC-39 is required for the synthesis of maradolipids but not of the trehalose headgroup, implying that OAC-39 most probably acts at the last step of maradolipid production, the acylation reaction.

### OAC-39 is enriched at the border to the lumen of the alimentary tube of dauer larvae

Identifying OAC-39 as an enzyme required for the last step of maradolipid synthesis opened the possibility to investigate where the maradolipid production occurs. Hence, we set out to study the localization of OAC-39. A protein reporter in which OAC-39 was translationally fused at the C-terimuns with eGFP was expressed in worms under the native *oac-39* promoter. Worms were then grown under high population density and the expression of OAC-39::eGFP was studied. As expected, OAC-39::eGFP fluorescence was only detected in dauer larvae and it showed a high association of OAC-39 with the gastrointestinal tract (Figure 2A). The highest expression was observed in the intestine. While it was visible as a reticulated or granulated pattern occupying the whole volume of the intestinal cells (Figure 2B, C, and D), it was much more concentrated at their apical surface, where it appeared to line the border to the gut lumen (Figure 2B). Markedly weaker expression was detected lining the pharyngeal and rectal cavities (Figure 2C and D). Thus, OAC-39 formed a continuum occupying the border to the lumen of the whole alimentary tube, consistent with the putative role of maradolipids in structuring the intestinal lumen and sealing the digestive system off from the external environment.

**Figure 2.**
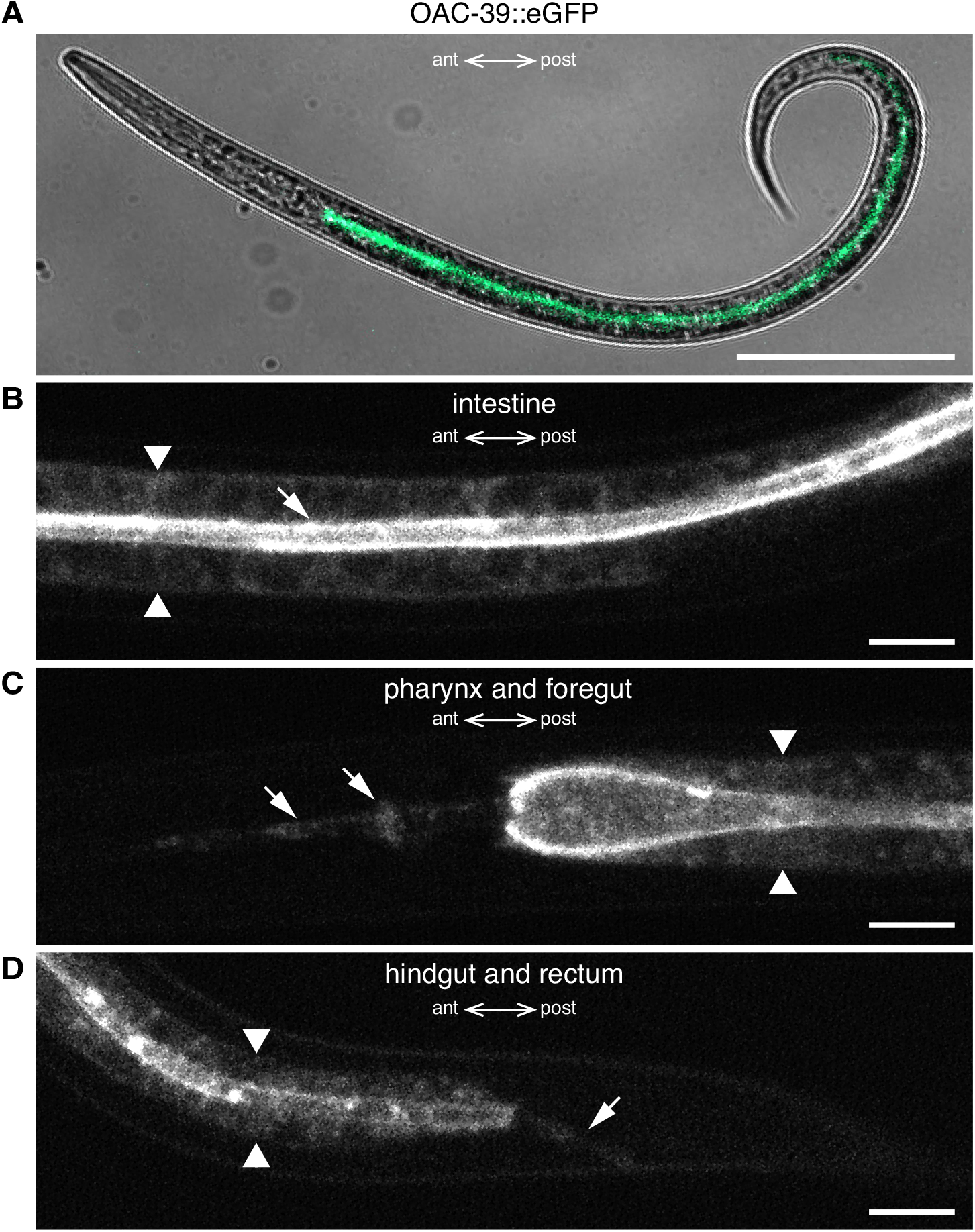
OAC-39 mainly localizes to the border of the lumen of the alimentary tube of dauer larvae. **A)** Overall expression of OAC-39::eGFP in a dauer larva. Composite bright-field (gray) and OAC-39::eGFP confocal fluorescence micrograph (green). Scale bar, 100 μm; ant – anterior; post – posterior. **B-D)** Confocal fluorescence micrographs of OAC-39::eGFP signal in different parts of the alimentary tract: B) in intestinal cells (arrowheads – basal sides, arrow – apical surface); **C)** in the pharynx and the foregut (arrowheads – basal sides of the foregut, arrows – lumen of the pharynx); D) in the hindgut and the rectum (arrowheads – basal sides of the hindgut, arrow – canal of the rectum). In **B-D)**, scale bars - 10 μm; ant – anterior; post – posterior.

## Discussion

In this study, we identified a dauer-specific *O*-acyltransferase, OAC-39, required for the synthesis of maradolipids and most probably also of lyso-maradolipids. The fact that OAC-39-deficient dauer larvae are depleted in maradolipids makes this enzyme the primary candidate for a “maradolipid synthase” incorporating acyl groups to trehalose. Although less likely, OAC-39 could indirectly stimulate the synthesis of trehalose lipids, e.g., by facilitating another catalytically active protein in performing the acyl transfer. Future studies employing orthologous expression of OAC-39 in organisms that lack trehalose lipids such as yeast or *in vitro* reactions will address its enzymatic activity.

Consistent with its role in lipid metabolism, OAC-39 has ten putative membrane-spanning domains and, therefore, must be mostly associated with cell membranes. In accordance, the reticulated/granulated intracellular localization of the reporter can be attributed to the presence of OAC-39 in the endomembrane system. The extensive apical enrichment of OAC-39, on the other hand, most probably results from its trafficking to the apical plasma membrane. Thus, it is highly probable that most of the acylation of trehalose that generates maradolipids and lyso-maradolipids occurs at the apical plasma membrane of intestinal cells. It remains to be determined if these trehalose lipids then remain incorporated in the plasma membrane, are released into the gut lumen, or internalized in the cells via endocytosis or some other transport mechanism. In the light of the putative role of maradolipids in forming the coat-like structure covering the luminal surface of the intestinal brush border, however, it is tempting to speculate that a significant portion of maradolipids remains apically presented.

The discovery of OAC-39 provides a valuable genetic and biochemical tool for investigating the function of trehalose lipids in dauer larvae. As bulk fraction of the dauer lipidome, (lyso-)maradolipids may act as structural lipids, surfactants, or modulate membranes’ physical-chemical properties, lipid droplets, and other lipophilic entities. On the other hand, they could be used as storage for trehalose and mmBCFAs. Because these compounds have been implicated in fat storage (32, 33), development (34–37), nutrient sensing signaling cascades (14, 36–38), and stress resistance (15, 16), (lyso-)maradolipids could affect some of these aspects of physiology through modulation of the availability of trehalose and mmBCFAs.

Finally, OAC-39 might be one of many enzymes in *C. elegans* that esterifies fatty acids to carbohydrates. Several other *oac* members display very high sequence similarity to *oac-39*. It is thus possible that some of them also produce trehalose lipids or similar glycosides of fatty acids at different locations or developmental settings. Studying the *oac* gene family in more detail will shed light on a novel metabolic module in *C. elegans* and other nematodes and perhaps bring new insights into the biochemical basis of developmental quiescence and stress resistance.

## Data availability

The data used during the current study are available from the author on reasonable request.

## Acknowledgments

Sider Penkov is grateful to Dr. Cihan Erkut and Prof. Teymuras Kurzchalia for sharing information on unpublished transcriptional analysis, to Prof. Hans-Joachim Knölker for the synthesis of lophanol, to Dr. Kathrin Schmeisser for critical reading, and to the Caenorhabditis Genetics Center, which is funded by NIH Office of Research Infrastructure Programs (P40 OD010440), for providing worm strains.

